# Enhancing single-cell western blotting sensitivity using diffusive analyte blotting and antibody conjugate amplification

**DOI:** 10.1101/2023.06.13.544857

**Authors:** Mariia Alibekova Long, William Benman, Nathan Petrikas, Lukasz J. Bugaj, Alex J. Hughes

## Abstract

While there are many techniques to achieve highly sensitive, multiplex detection of RNA and DNA from single cells, detecting protein contents often suffers from low limits of detection and throughput. Miniaturized, high-sensitivity western blots on single cells (scWesterns) are attractive since they do not require advanced instrumentation. By physically separating analytes, scWesterns also uniquely mitigate limitations to target protein multiplexing posed by affinity reagent performance. However, a fundamental limitation of scWesterns is their limited sensitivity for detecting low-abundance proteins, which arises from transport barriers posed by the separation gel against detection species. Here we address sensitivity by decoupling the electrophoretic separation medium from the detection medium. We transfer scWestern separations to a nitrocellulose blotting medium with distinct mass transfer advantages over traditional in-gel probing, yielding a 5.9-fold improvement in limit of detection. We next amplify probing of blotted proteins with enzyme-antibody conjugates which are incompatible with traditional in-gel probing to achieve further improvement in the limit of detection to 10^3^ molecules, a 520-fold improvement. This enables us to detect 85% and 100% of cells in an EGFP-expressing population using fluorescently tagged and enzyme-conjugated antibodies respectively, compared to 47% of cells using in-gel detection. These results suggest compatibility of nitrocellulose-immobilized scWesterns with a variety of affinity reagents — not previously accessible for in-gel use — for further signal amplification and detection of low abundance targets.

## Introduction

Variability among single cells (heterogeneity) is a fundamental property of cell populations. Cell functional states change rapidly, especially during development and differentiation ^1,2^. Changes in cell function affect nearby cells, leading to development of heterogeneous tissues from initially identical cells ^2–4^. Disease is also characterized by changes in cell state heterogeneity that affects its progression and complicates development of disease models and treatments ^3,5–7^. Proteins and their post-translational modifications are known to regulate cell functional states more proximally than mRNA and thus provide more information about cell fate decisions than single-cell transcriptomics ^8,9^.

However, quantifying single cell proteome states remains a significant technical challenge in large part because proteins and PTMs — unlike mRNA — cannot be amplified to aid detection. Bridging this capability gap between single-cell proteomics and transcriptomics would enable higher-resolution functional mapping of single cells.

Two main groups of methods are currently employed in single-cell proteomics, including highly sensitive antibody-based methods and highly-multiplexed mass spectrometry ^10–17^. Among antibody-based methods, microfluidic assays that run on open-faced hydrogels have significant advantages in simplicity and modularity, enabling several analytical advances^18–20^. Hydrogel matrices can sieve, concentrate, or immobilize proteins, enabling quantitation of species in downstream assay steps. However, hydrogel properties favorable for one step are often not favorable for others and can create conflicts of performance between them. For example, in single-cell western blotting (scWesterns), cells are loaded into microwells stippled into a thin polyacrylamide (PA) gel, lysed, and their protein contents separated by electrophoresis^18^. Proteins are then covalently photocaptured in the gel using photoactive benzophenone-methacrylamide (BPMAC) co-monomers and probed using fluorescently-labeled antibodies^21,22^. The protein capture step fixes analytes in place, avoiding a rapid erosion of spatial resolution due to diffusion. However, this integration of separation and in-gel detection creates mass-transfer conflicts between analyte sieving during separation, which requires gel pore sizes on the order of protein sizes, and in-gel probing, which favors larger pore sizes to increase the penetration of antibodies^18,23^. Small pore-size polyacrylamide gels tend to prevent antibody penetration due to partitioning, which limits their in-gel concentration at equilibrium^23,24^. This problem is exacerbated when using bulkier detection reagents such as antibody-enzyme conjugates that could otherwise add sensitivity or analyte multiplexing benefits. Even for fluorescently-labeled antibodies, ∼10-fold higher free-solution concentrations are required compared to traditional membrane immunoblotting to achieve sufficient in-gel concentrations^18^. This in turn can result in relatively high background staining of gels due to physical trapping of unbound antibody, limiting assay performance and increasing cost^25^.

Diffusional resistance can be circumvented using out-of-plane electrotransfer of antibodies^26^ or chemical modification of gel pore size (including between assay steps)^27,28^. However, each of these require significant increases in assay complexity and likely still limit compatibility of scWesterns with alternative detection schemes. Therefore, significant potential remains to circumvent in-gel probing in scWesterns while retaining the ability to ‘lock in’ spatial resolution of separated proteins on the microscale.

Here we replace in-gel probing with an alternative diffusive transfer step of analytes to a large pore-size nitrocellulose membrane better suited for antibody probing (**Figure 1**). This not only significantly increases antibody probing sensitivity but also allows the use of enzyme-antibody conjugates for amplified target detection, which has not been possible in polyacrylamide gels. We find that nitrocellulose blotting of scWestern separations operates at >90% transfer efficiency for a wide range of protein molecular weights (30-150 kDa) and in an immobilization-dominated regime relative to diffusion. This better limits band spreading during blotting relative to traditional in-gel scWesterns. We find a 5.9-fold improvement in the limit of detection for EGFP quantitation on paper relative to in-gel quantitation using primary and fluorescently-labeled secondary antibodies. Additionally, we used a 10-fold lower antibody concentration, thus reducing non-specific background and assay cost. Leveraging the relatively accessible pore sizes of nitrocellulose that allows for bulkier detection species, we assayed substrate turnover from enzyme-linked antibodies to achieve 520-fold improvements in limit of detection, down to 10^3^ molecules. These improvements were sufficient to detect a > two-fold wider range of expression heterogeneity in EGFP from HEK293 cells than achievable in polyacrylamide gels.

**Figure 1:**
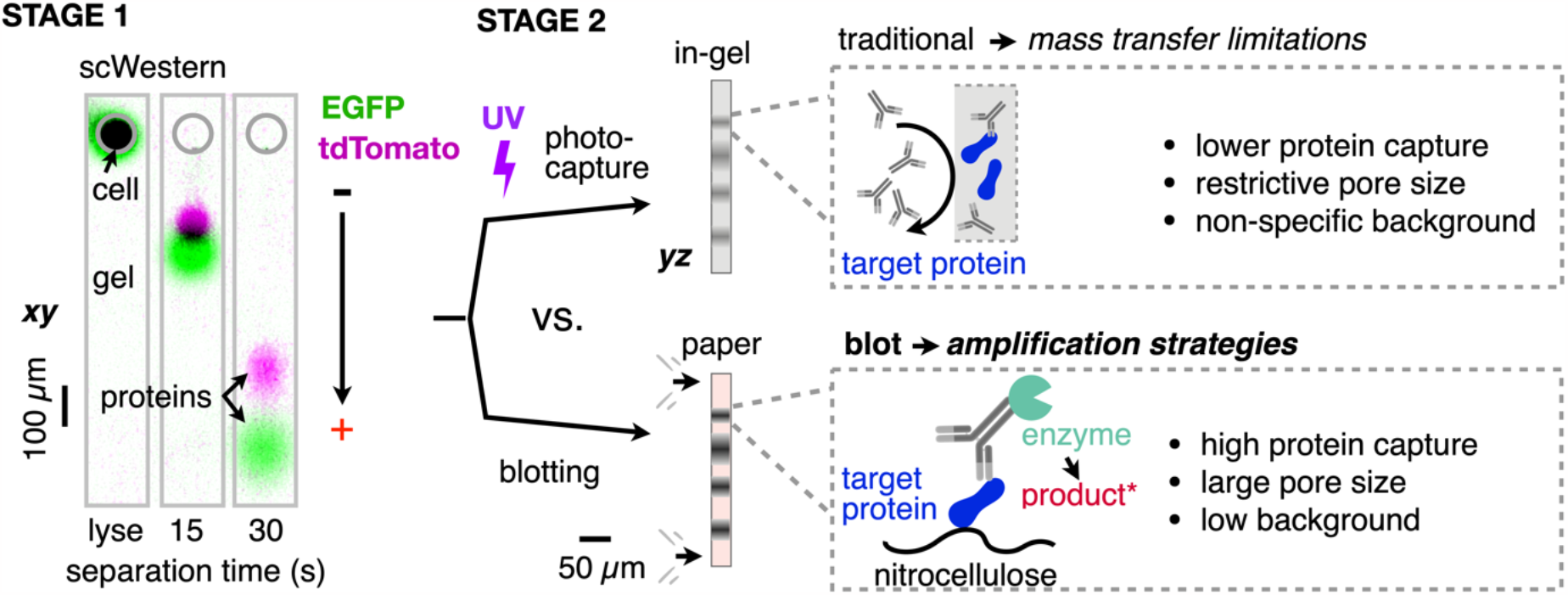
Diffusive blotting to porous nitrocellulose paper for single-cell western blotting with enhanced sensitivity. Stage 1 - scWesterns separate proteins from polyacrylamide microwell-localized single cells or purified proteins by non-reducing SDS-PAGE (here EGFP, 27 kDa; tdTomato, 55 kDa from a microwell containing a single lysed HEK293 cell). Stage 2 - In traditional scWesterns, analytes are photocaptured inside the sieving gel. Here we instead perform blotting to nitrocellulose paper by diffusion.

## Experimental Section

### Protein immobilization on nitrocellulose membranes and protein transfer efficiency

1.5 μL of 5 μM GFP (27 kDa), Ovalbumin-AlexaFluor 555 (46 kDa), or IgG-AlexaFluor 555 (150 kDa) were spotted on a 7%T (total monomer) 3.3%C (weight % bisacrylamide crosslinker relative to total monomer) PA gel ^18,22,29^, incubated for 5 min at room temperature and transferred upon contact with 0.2 μm pore-size nitrocellulose for 1-4 min. Fluorescence of protein spots on gel pre- and post-transfer was compared to determine transfer efficiency. See *Supporting Information* for detail.

### Diffusive spreading of immobilized protein

Diffusive spreading of single-cell proteins spatially separated by single-cell SDS-PAGE and immobilized on 1) BPMAC-functionalized polyacrylamide or 2) nitrocellulose via transfer from plain polyacrylamide gel for 1-4 min was quantified by increase in spot size. See *Supplementary Information* for detail.

### Microwell concentration calibration

PDMS microchannels were fabricated with channel height 30 μm (identical to polyacrylamide gel thickness) and adhered to glass slides (see *Supplementary Materials*). 10 μL of GFP-Alexa 555 at concentrations of 0.001, 0.01, 0.1, 1, 5, and 10 μM were added to microchannels and imaged on a confocal microscope to generate a calibration curve. Fluorescence was plotted against EGFP molecule number for calibration and quantification of limit of detection.

### Protein partition coefficient measurement

40 μL of 0.1 μM or 1 μM of EGFP-Alexa Fluor-555 in RIPA-like buffer (0.5% SDS, 0.1% v/v Triton X-100, 0.25% sodium deoxycholate in 12.5 mM Tris, 96 mM glycine, pH 8.3, 0.5× from a 10× stock) were added to a BPMAC-functionalized polyacrylamide gel with 20 μm-diameter wells, sandwiched in a hybridization cassette. The gel was then briefly rinsed, sandwiched with a glass slide, and EGFP fluorescence was measured in gel and in microwells on a confocal microscope and correlated. The partition coefficient was calculated from the slope of the best fit line and used to determine in-gel and in-well concentrations relative to EGFP solution concentrations.

### Fluorescently-labeled antibody probing calibration

On 1) scWestern gel; 0.5 μL of 0.01, 0.0193, 0.0373, 0.072, 0.1389, 0.2683, 0.5179, and 1 μM solutions of EGFP-AlexaFluor 555 in RIPA-like buffer were spotted and fixed by irradiating with 254 nm UV light (7 mW cm^-2^) for 60 s. Antibody probing with either 1:200 goat anti-GFP or 1:400 rabbit polyclonal anti-GFP antibody both at 0.05 mg ml^-1^ for 2 hr was then performed at room temperature and appropriate secondary antibody-AlexaFluor 647 at 1:500 dilution at 0.004 mg ml^-1^ for 1 hr at room temperature. On 2) nitrocellulose, 0.5 μL of the identical protein concentrations were spotted on scWestern gel and transferred to nitrocellulose for 3 min. The membrane was fixed, blocked and probed with antibodies as described above. See *Supplementary Materials* for details.

### Enzyme-linked antibody probing calibration

0.5 μL of 1 μM, 100 nM, 10 nM, 1 nM, 100 pM, 10 pM, 1 pM, 100 fM, and 0 μM EGFP in RIPA-like buffer were spotted on a 7% PA gel and transferred to nitrocellulose for 3 min. The membrane was fixed, blocked and probed with antibody as described above except secondary probing was with 1:10,000 dilution of horseradish peroxidase (HRP)-conjugated rabbit anti-goat secondary antibody. We then cut nitrocellulose with protein blots and placed them face down in 48-well microplate wells. Detection was performed via QuantaRed Chemifluorescence kit (ThermoFisher Scientific 15159) to convert ADHP substrate to resorufin in the presence of HRP and H_2_O_2_ applied as a 100 μL reaction cocktail per well. Resorufin fluorescence was detected by widefield microscopy. Slopes of fluorescence vs. time plots were used to create a calibration curve. LOD was determined as the lowest analyte concentration at which the fluorescence increase was greater than the increase in standard deviation of the blank in a reaction time equal to the time *t*_*D*_ that resorufin diffuses a characteristic distance of 100 μM (the approximate size of single-cell protein plumes in scWestern separations). See *Supplementary Materials* for details.

### Single-cell primary and secondary antibody-based and enzyme-linked antibody protein detection

We tested and compared 1) standard BPMAC in-gel immobilization and probing with primary and secondary fluorescently tagged antibody vs 2) the same for nitrocellulose blotting, 3) enzyme-linked antibody-based EGFP detected in scWestern blots bearing single-cell separations. For 1) <u>BPMAC-immobilized</u> single-cell proteins, gel was incubated with 1:20 polyclonal goat anti-GFP antibody for 2 hr at room temperature, washed and incubated with 1:50 dilution of donkey anti-goat Alexa-555 antibody in 1% BSA in TBST for 60 min at room temperature. For 2) <u>nitrocellulose-based detection</u>, nitrocellulose was probed with 1:200 primary antibody for 2 hr at room temperature, washed and incubated with 1:500 donkey anti-goat Alexa-555 antibody for 60 min at room temperature. For 3) <u>Enzyme-linked detection on nitrocellulose</u> in scWesterns, primary antibody probing was performed with 1:200 goat anti-GFP for 2 hr at room temperature, washed and probed with rabbit anti-goat secondary IgG antibody conjugated to HRP for 60 min at room temperature. 100 μL of QuantaRed kit was added to nitrocellulose bearing single-cell Western protein separations, sandwiched with a glass slide and signal imaged on a widefield microscope. We determined background-adjusted average fluorescence values of single cell EGFP spots immediately after immobilization. Next, we found background adjusted average fluorescence of corresponding ROI readouts for 1) BPMAC-immobilized protein secondary antibody signal, 2) nitrocellulose-immobilized secondary antibody signal and 3) enzyme-linked antibody resorufin signal increase. We defined LOD as 3.3 times standard deviation of background for 1) and 2) and 3.3 times standard deviation of blank sample at *t*_*D*_ for 3) enzyme-linked detection. We then plotted initial EGFP fluorescence values to corresponding readout values and determined % of cells below LOD. See *Supplementary Materials* for details.

## Results and Discussion

### Transfer of proteins from polyacrylamide gel to nitrocellulose is efficient and retains spatial resolution

Previous scWestern implementations noted significant sensitivity barriers for in-gel antibody probing, since IgG species suffer from a 5.9x lower equilibrium concentration in 8%T polyacrylamide relative to free solution due to partitioning ^18^. Additionally, in-gel antibody diffusion required equilibration time of 4*τ* ∼ 3 min during each probing and washing step ^18^. Diffusive transfer of protein species from polyacrylamide gels to blotting media has been reported for traditional separation assays^30–32^. However, its performance has not been validated at spatial scales relevant to scWesterns. We reasoned that transferring protein species from single-cell separations to a nitrocellulose paper interface^33–36^ would increase sensitivity by mitigating subsequent mass transfer limitations during probing. It would also enable alternative assay readout strategies such as enzyme-linked antibody detection that is not compatible with the restrictive pore sizes of polyacrylamide gels. Transfer to nitrocellulose paper would also simplify the scWestern workflow by allowing the use of traditional polyacrylamide gel formulations rather than those with photoactive capture functionality. Finally, analyte detection could achieve higher signal-to-noise ratios on-paper due to relatively low non-specific antibody binding ^37,38^.

We began by performing scWestern separations of fluorescent EGFP and tdTomato proteins expressed in HEK 293T cells, finding that proteins appeared to blot successfully after immediately interfacing the separation gel with nitrocellulose (**Figure 2A,B**). Two performance parameters are crucial to high-fidelity blotting -transfer efficiency and spatial resolution. We first sought to evaluate transfer efficiency for purified proteins embedded in scWestern gels and diffusively transferred to a 0.2 μm pore size nitrocellulose membrane for 1-4 min (see Experimental Section). This time range bracketed a theoretical transfer time τ ∼ L^2^/D for a characteristic diffusion length equal to gel thickness (30 μm) and EGFP diffusivity of ∼38 μm^2^ s^-1^ in 7%T 3.3%C PA gel, such that τ ∼ 24 s and 5τ ∼ 2 min ^22,23,39–41^. Next, we imaged the gel for residual protein and quantified transfer efficiency as:

**Fig. 2:**
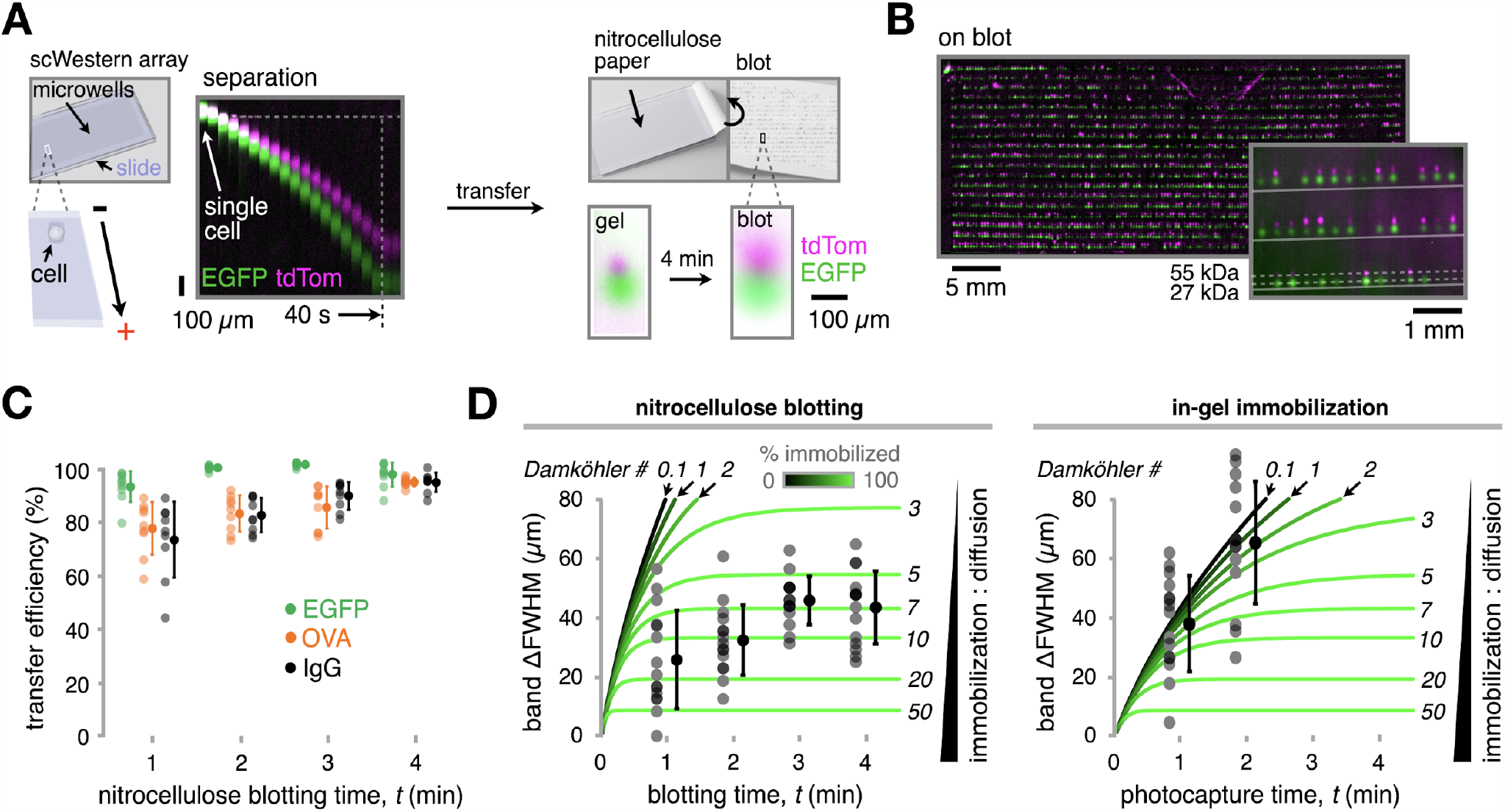
Nitrocellulose blotting achieves high transfer efficiency and improved band dispersion compared to traditional scWesterns. (**A**) Schematic and fluorescence micrographs of single-cell protein separation and blotting to nitrocellulose. (**B**) Fluorescence micrographs showing single-cell protein separations post-blotting. (**C**) Plot of protein spot transfer efficiency from gel to nitrocellulose paper for varying blotting times. (**D**) Loss in band resolution determined by increase in full-width at half-maximum (ΔFWHM) for pre-vs. post-blotted EGFP bands after single-cell separation. Experiment data are overlaid on simulation trajectories for different Damköhler numbers showing expected band dispersion and % of EGFP molecules immobilized over time.

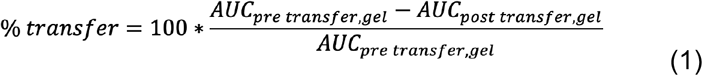

where AUC is background-subtracted fluorescence area under the curve for the ROI. Transfer efficiency exceeded 90% for ∼150 kDa and smaller proteins after 4 min, with even the heavier IgG marker exceeding 80% efficiency after 2 min (**Figure 2C**). These data indicate efficient analyte efflux from the gel during nitrocellulose blotting for the studied range of protein molecular mass. This range spans the median protein molecular weight of ∼50 kDa ^42^, predicting highly efficient transfer for many single-cell protein targets using the nitrocellulose blotting approach.

One concern for microscale scWestern blotting relative to traditional blotting is that the rate of analyte diffusion may become significant relative to the rate of immobilization to the blotting medium at very small characteristic lengths. This would decrease separation resolution, reduce concentrations of target protein available for analysis due to diffusive band spreading, and negatively affect assay limit of detection. We therefore sought to quantify band spreading during transfer of EGFP bands from single-cell separations to nitrocellulose over a range of blotting durations (**Figure 2D**). The increase in full-width-at-half-maximum (FWHM) values was modest over the blotting time — generally less than 50 μm for initial EGFP band widths of ∼100 μm. Spreading appeared to saturate after 3 min, implying that analyte immobilization was complete in that time. The kinetics of band spreading are a function of a Damköhler number^22,43,44^ (*Da = kL*^*2*^*/D*), which represents the ratio of the characteristic timescales of diffusion (*L*^*2*^*/D*) and immobilization (*1/k*) where *k* is a first-order immobilization rate constant (s^-1^), *L* is a characteristic diffusion length that we take to be the initial band width (FWHM), and *D* is diffusivity of EGFP in nitrocellulose (we assume similar to that in free solution at ∼90 μm^2^ s^-1^, ref. ^22,39–41^, see *Supplementary Methods*). To more quantitatively understand if band spreading is diffusion-or immobilization-dominated during blotting, we simulated 1D diffusion and immobilization of a Gaussian band of EGFP in paper (initial FWHM of 100 μm), ignoring mass transfer resistance from gel to paper:

For the free solution fraction:

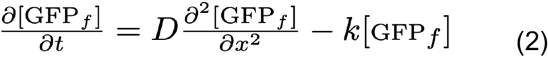

For the immobilized fraction:

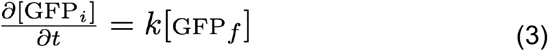

Where the subscripts *f* and *i* denote ‘free-solution’ and ‘immobilized’, respectively. Since the immobilization rate constant *k* is unknown, we simulated a range of values creating a corresponding range of Damköhler number values, where *Da* < 1 indicates kinetics are diffusion-dominated, and *Da* > 1 indicates kinetics are immobilization-dominated. We found that the EGFP single cell transfer data fell within a Damköhler number range of ∼5-10 from the model, predicting favorable immobilization-dominated kinetics. In practice, we also fixed transferred proteins to the blotting medium using paraformaldehyde to preserve high transfer efficiency and protein spot bandwidth. We then performed a similar analysis comparing nitrocellulose blotting against BPMAC-mediated in-gel immobilization in traditional scWesterns (**Figure 2D**). In contrast to nitrocellulose blotting, transport appeared to be far more diffusion-dominated for BPMAC-mediated immobilization as indicated by higher band spreading. Sufficient sensitivity to analyze the spatial distribution of the bound EGFP fraction was only possible at 1 and 2 minutes due to band spreading and, potentially, UV-induced bleaching. To evaluate diffusion and immobilization kinetics in BPMAC-mediated protein capture, we created a second version of the model, where *D* was the diffusivity of EGFP in a 7%T scWestern gel (∼38 μm^2^ s^-1^, ref. ^22,39–41^). Despite its lower diffusivity in-gel, EGFP immobilization fell within a Damköhler number range of ∼0.1-3, predicting greater diffusive band spreading relative to immobilization compared to nitrocellulose blotting. These data validate rapid and efficient immobilization of microscale protein bands to the nitrocellulose blotting medium, retaining spatial resolution in scWestern separations.

### Nitrocellulose blotting increases detection sensitivity and reduces antibody use compared to in-gel immobilization

Having validated favorable blotting performance, we sought to compare analyte detection sensitivity for in-gel probing vs. probing on nitrocellulose blots (**Figure 3A**). Direct comparison between the two methods for single cells is not possible since fluorescent fusion protein expression is heterogeneous in cell cultures. Instead, we mimicked immobilization from single cells in BPMAC gels vs. after transfer to nitrocellulose using 0.5 μl spots of a serial dilution of purified AlexaFluor 555-labeled EGFP. We labeled EGFP to increase its capture efficiency by BPMAC^22^ to make it as competitive as possible with nitrocellulose blotting. For BPMAC-based immobilization, equilibrated spots were photo-immobilized in BPMAC polyacrylamide scWestern gels through a photomask, antibody probed, and imaged. We then determined spot signal-to-noise ratio (SNR, see *Supplementary Information*) and plotted it against number of EGFP molecules determined by estimating in-gel [EGFP] using its partition coefficient (**Figure S1**) and taking a gel volume equivalent to that of a ∼100 μm diameter single-cell spot. For nitrocellulose-based immobilization, spots equilibrated in plain polyacrylamide scWestern gels were transferred from the gel to nitrocellulose for 3 minutes before performing a similar analysis. Primary and fluorescently-labeled secondary antibody concentrations were optimized for the lower concentration range typical of traditional nitrocellulose blotting rather than the substantially higher concentrations (∼10-100-fold) used in in-gel scWesterns^18^. We found an optical limit of detection (LOD) of ∼10^6.5^ molecules of EGFP for traditional in-gel detection, and ∼10^5.7^ molecules for detection on nitrocellulose; a 5.9-fold improvement in detection sensitivity using nitrocellulose blotting (see *Supplementary Methods* for calculation details). With a second primary anti-GFP antibody we obtained a similar LOD of 10^6.9^ EGFP molecules for traditional in-gel detection and 10^6^ molecules for nitrocellulose detection (7-fold improvement) (**Figure S2A**). These data reveal significant improvement in detection sensitivity for EGFP blotted to nitrocellulose relative to photocapture and probing in-gel.

**Figure 3:**
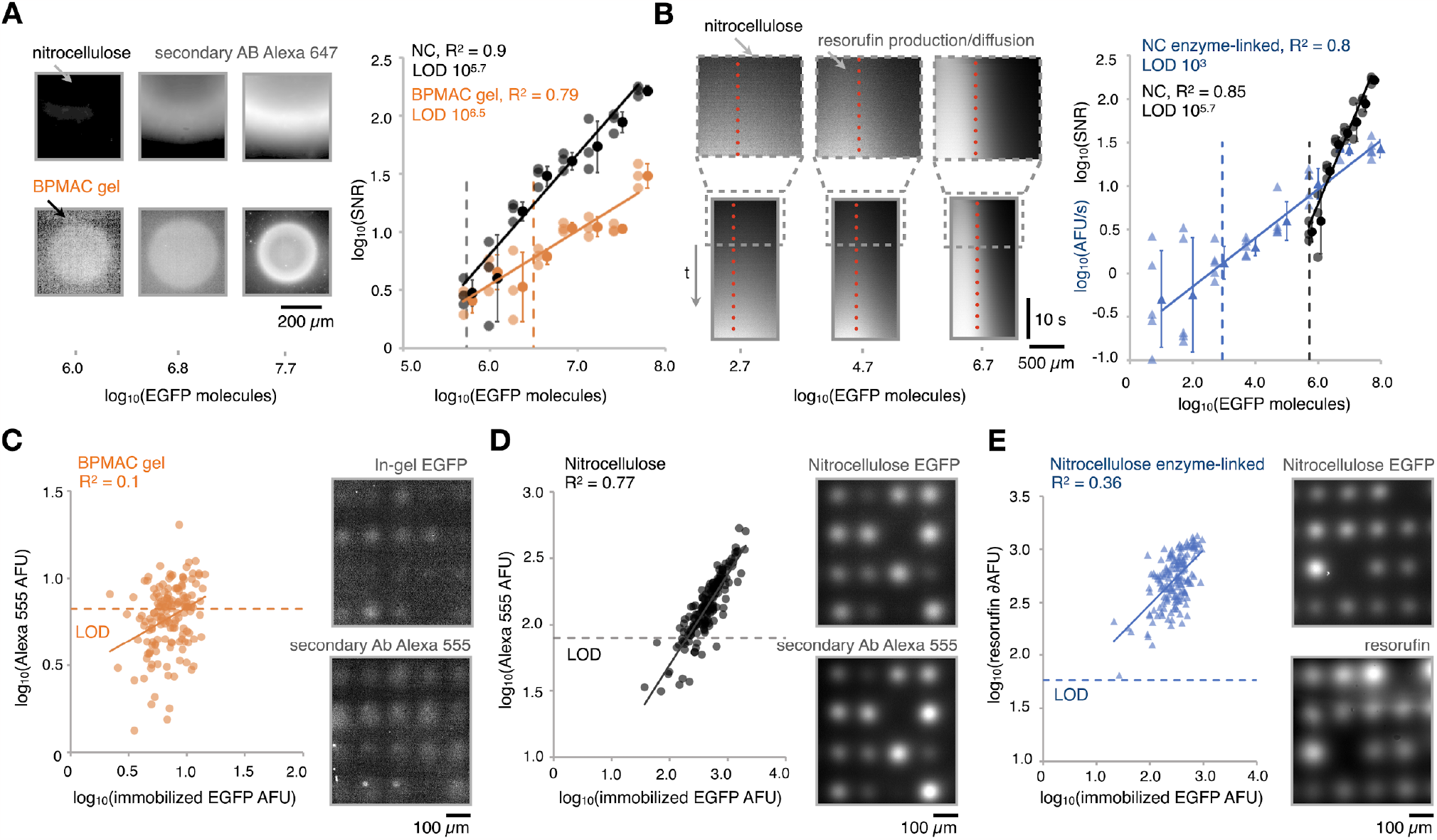
Nitrocellulose blotting increases assay sensitivity and enables enzyme-based amplification of scWestern readouts. (**A**) *Left*, immunofluorescence micrographs of calibration spots of EGFP transferred to nitrocellulose or immobilized in-gel prior to probing with goat anti-EGFP primary and anti-goat Alexa-647 secondary antibody. Pixel intensity was log-transformed. Note that contrast settings are equivalent within each image row, but different between rows. *Right*, corresponding calibration curves. (**B**) *Left*, immunofluorescence kymographs of calibration spots of EGFP transferred to nitrocellulose and detected with HRP-conjugated secondary antibodies for enzymatic amplification. Dotted line shows initial protein spot boundary before resorufin diffusion. Inset shows kymographs for the first 10 s. *Right*, corresponding calibration curve compared to fluorescently-labeled secondary antibody detection. (**C**) *Left*, intensity of secondary antibody signal detecting EGFP bands vs initial immobilized EGFP intensity, log transformed, from single-cell separations on BPMAC-immobilized polyacrylamide gel. *Top right*, EGFP fluorescence micrograph immediately post immobilization in BPMAC gel. *Bottom right*, secondary antibody-Alexa 555 fluorescence micrograph for the same ROI. (**D**) *Left*, intensity of secondary antibody signal detecting EGFP bands vs initial immobilized EGFP intensity, log transformed, from single-cell separations on nitrocellulose. *Top right*, EGFP fluorescence micrograph immediately post immobilization in nitrocellulose. *Bottom right*, secondary antibody-Alexa 555 fluorescence micrograph for the same ROI. (**E**) *Left*, resorufin signal increase over *t<t*_*D*_ detecting EGFP bands vs initial immobilized EGFP intensity, log transformed, from single-cell separations on nitrocellulose. *Top right*, EGFP fluorescence micrograph immediately post immobilization in nitrocellulose. *Bottom right*, resorufin fluorescence micrograph for the same ROI at *t* = *t*_*D*_.

### Nitrocellulose blotting enables 520-fold improvement in sensitivity via enzyme-linked antibody detection

Free from the limitations of in-gel probing, we next attempted to improve detection sensitivity. We employed HRP-conjugated secondary antibody and assayed for ADHP substrate (10-acetyl-3,7-dihydroxyphenoxazine) conversion to fluorescent resorufin in the presence of H_2_O_2_ to read out the assay directly on nitrocellulose paper bearing individual EGFP protein spots. The rate of resorufin fluorescence increase here is expected to be proportional to the local HRP concentration, given excess ADHP ^45–49^. Calibration showed a dynamic range of ∼10^3^-10^8^ molecules of EGFP (**Figure 3B**). Since the resorufin product is soluble, analyte detection is a race between resorufin production and diffusion, with characteristic times *t*_*R*_ and *t*_*D*_ respectively. In other words, successful analyte detection while retaining acceptable spatial resolution would only occur when *t*_*R*_ < *t*_*D*_. Across the assay dynamic range, *t*_*R*_ (time for resorufin signal to exceed 3.3*S.D. of background) was smaller than *t*_*D*_ = 10.4 s estimated for resorufin to diffuse an equivalent distance to a ∼100 μm band in scWestern applications (see *Supplementary Information*). These data show that enzyme-linked detection strategies yield favorable performance in scWestern assays down to a detection limit of ∼10^3^ molecules of EGFP without significant loss in protein band resolution. This yields ∼520 fold improvement in LOD over fluorescently-tagged antibody detection, enabling future application to detection of low abundance proteins. Additionally, we verified that enzyme-linked detection failed in scWestern gels (**Figure S2B**). Nitrocellulose blotting of single-cell protein separations may therefore enable a variety of amplification strategies that are incompatible with in-gel detection.

### Nitrocellulose blotting improves measurement of single-cell protein expression heterogeneity

To test the performance of our diffusive analyte blotting approach for single-cell protein detection, we assayed for EGFP in scWestern blots using fluorescently-tagged antibody detection and the enzyme-antibody amplification scheme (**Figure 3C-E**). We detected only 46.6% of EGFP+ cells previously FACS sorted for high EGFP expression in BPMAC-immobilized blots (70 of 150 cells). On nitrocellulose blots, assaying the same cell culture using fluorescently-labeled secondary antibody successfully detected EGFP in 86.5% of cells (138 of 160 cells). Finally, EGFP was detected in 100% of the cells assayed using enzyme-linked antibody amplification on nitrocellulose (160 of 160 cells). We note that the observed positive linear correlation between resorufin fluorescence increase and initial fluorescence is weaker than that detected by fluorescently-tagged antibodies on nitrocellulose (R^2^ of 0.36 vs 0.77 respectively). We hypothesize that ADHP mass transfer limitations may contribute to this additional variability given that ADHP may become limiting in areas of scWestern blots with locally denser separations. While enzyme-antibody conjugate-based detection of single-cell proteins is highly sensitive here, further innovation may be needed to remove mass transfer limitations on local substrate concentration to increase assay linearity and quantitative capability. Overall, our data demonstrate successful application of alternative antibody conjugate-based detection in scWesterns. This innovation improved detection sensitivity and may offer advantages in analyte multiplexing over in-gel probing in future work.

## Conclusions

We introduce diffusive scWestern blotting to nitrocellulose for microscale protein bands, improving assay sensitivity and adaptability to different detection schemes relative to traditional in-gel antibody probing. We found that nitrocellulose blotting has favorable analyte retention and band spreading characteristics, validated by reaction-diffusion modeling. Head-to-head comparison of detection limits for fluorescently-labeled antibody detection of EGFP revealed a 5.9-fold improvement using on-paper analyte blotting relative to in-gel photocapture. Nitrocellulose blotting allowed us to use 10-fold lower antibody concentrations due to the lack of antibody probe partitioning that limits in-gel immobilization. This significantly reduces background, increases signal-to-noise ratio and improves characterization of single-cell expression heterogeneity. Additionally, nitrocellulose-based immobilization significantly reduces assay complexity.

Limits of detection improved to as few as 10^3^ molecules using enzyme-antibody conjugates for detection, a 520-fold improvement in sensitivity. This amplification strategy had previously not been compatible with in-gel scWesterns due to the prohibitive pore-size of the combined separation/blotting medium. Transferring separations to a large pore-size medium may enable future application of a larger diversity of amplification strategies for *in situ* detection beyond those applied here, including chemiluminescence, rolling circle amplification, hybridization chain reaction, nanoparticle/quantum dot-based techniques etc. These may also ease mating of scWesterns to other detection approaches such as mass spectrometry^16^. Future work will aim to complement these advances to improve scWestern multiplexing, which would have transformative impacts in single-cell proteomics.

## Acknowledgements

The authors thank members of the Hughes and Bugaj lab for discussions and support. This work was supported by the National Institutes of Health (NIGMS R21GM132831 to A.J.H. and L.J.B.). M.A.L. was supported by an American Society of Nephrology and KidneyCure Pre-Doctoral Fellowship and W.B. was supported by an NSF Graduate Research Fellowship.

## Supporting Information

### Reagents and materials

All following chemicals and materials were received from manufacturers. Glass slides (Corning 2947-75X50), nitrocellulose membrane 0.2 μm pore size (ThermoFisher Scientific, 77012), silicone wafers (University Wafer), SU8 2025 photoresist (Y111069, MicroChem), SU8 developer (ThermoFisher Scientific 14200-075), dichloromethylsilane (DCDMS, Sigma Aldrich 440272), acrylamide/Bis-acrylamide 30% solution 37.5:1 (Sigma-Aldrich, A3699-100ML), BPMAC (PharmAgra),SDS (Bio-Rad, 161-0301), Triton X-100 (Sigma Aldrich, T9284-1L), ammonium persulfate (APS, Sigma-Aldrich, A3678), tetramethylethylenediamine (TEMED, Sigma-Aldrich, T9281), 10X TBS (BioRad, 1706435), 10X DPBS(ThermoFisher Scientific 14200-075), EDTA (Sigma-Aldrich, 03690-100ML), polyoxyethylene (20) sorbitan monolaurate (similar to Tween-20, OmniPur, 9480),QuantaRed kit chemifluorescence kit (ThermoFisher Scientific, 15159), Alexa-555 (ThermoFisher Scientific, A20009). All following proteins and antibodies were received from manufacturers. Ovalbumin Alexa Fluor-555 conjugate (ThermoFisher Scientific, O34782), bovine serum albumin (Sigma-Aldrich, A2153-100G), goat anti-GFP polyclonal antibody (Abcam, ab6673), rabbit anti-GFP polyclonal antibody (Thermo Scientific, A-11122), donkey anti-rabbit Alexa Fluor-647 secondary antibody (ThermoFisher Scientific, A-31573), donkey anti-goat Alexa Fluor-647 secondary antibody (ThermoFisher Scientific, A-32849), rabbit anti-goat HRP conjugated secondary antibody (ThermoFisher Scientific, A-31402). Following buffers were in-house made: RIPA-like lysis buffer (2.5 mg/mL Sodium Deoxycholate, 5 mg/mL SDS, 0.1% Triton X-100, 0.5X Tris-Glycine), 1X TBST (1X TBS, 0.1% Tween-20).

#### Cell lines and culture

HEK 293T cells were a gift from Lukasz Bugaj (UPenn, Takarabio Cat # 632180). HEK-293 cells were maintained in DMEM culture media (ThermoFisher Scientific, 11965084) with 10% FBS (Corning, 35-010-CV) and 1x Pen/Strep (Mediatech, MT30-002-CI). Cells were cultured in polystyrene flasks and kept in a humidified environment at 37ºC and 5%CO_2_.

#### Lentiviral packaging and cell line generation

Lentivirus was packaged by cotransfecting the pHR transfer vector, pCMV-dR8.91 (Addgene, catalog number 12263), and pMD2.G (Addgene, catalog number 12259) into Lenti-X HEK 293 T cells. Briefly, cells were seeded one day before transfection at a density of 350,000 cells ml_*−*1_ in a six-well plate in 2mL total. Plasmids were transfected using the calcium phosphate method. Media was removed one day post transfection and replaced with fresh media. Two days post transfection, media containing virus was collected and centrifuged at 800g for 3 min. The supernatant was passed through a 0.45-μm filter. 500 μl of filtered virus solution was added to 700,000 untransfected HEK 293 T cells seeded in a six-well plate. Cells were expanded over multiple passages, and successfully transduced cells were enriched through fluorescence-activated cell sorting (Aria Fusion).

#### Plasmid Assembly

GFP and tdTomato viral plasmids were generated by digesting the PHR transfer vector with MluI-HF and NotI-HF (New England Biolabs). GFP and tdTomato coding DNA fragments were generated via PCR and inserted into the digested backbone via HiFi cloning mix (New England Biolabs).

#### Plasmid transfection

HEK 293 T cells were transfected using the following calcium phosphate method: 150 μl of 2× HEPES-buffered saline (HeBS), 1.5 μg of each DNA construct and H_2_O up to 132 μl was added per 2 ml of media of the cell culture to be transfected. 18 μl of 2.5 mM CaCl_2_ per 2mL of media was added to the transfection mix after mixing of the initial components, incubated for 1 min 45 s at room temperature and added directly to the cell culture.

#### Fabrication of polyacrylamide gels

30 μm thick polyacrylamide gels were fabricated on methacrylate-functionalized glass slides against micropost arrays bounded by SU-8 shims adhered to silicon wafers as described previously^18^. Briefly, we spun SU-8 2025 photoresist (Y111069, MicroChem) onto mechanical grade silicon wafers (University Wafer) and baked according to manufacturer’s specifications. Photoresist was then exposed to 365 nm UV light (ThorLabs LED M365LP1, lens ACL7560U, lens tube SM3V10) through 20,000 dpi mylar mask (ArtNet Pro) to achieve a total 150 mJ/cm^2^ exposure. The mask was designed to include shims on the sides of the glass slide to allow for a 30 μm gap between wafer and glass slide to control thickness of fabricated polyacrylamide gel. The mask also contained a pattern to allow for generation of posts on wafer post-exposure, that will in turn allow us to fabricate microwells in the polyacrylamide gel. Wafers were processed with SU-8 developer solution (ThermoFisher Scientific 14200-075) with alternating acetone and isopropanol rinse. Wafers were silanized with dichloromethylsilane (DCDMS, Sigma Aldrich 440272) for 20 minutes *in vacuo*, rinsed with DI water and dried under a stream of compressed air.

Wafers could then be reused multiple times and cleaned with 0.1% Triton X 100 and DI water between each use. Gels were then cast on glass slides (Corning 2947-75X50) silanized with methacrylate functional groups as previously described ^18^. The gel precursor solution were 7%T (w/v total acrylamides), 2.7%C (w/w of cross-linker N,N-methylenebisacrylamide) from pre-mixed stock solution (A3699, Sigma-Aldrich), 10% 10x DPBS (ThermoFisher Scientific 14200-075), 0.05 % SDS (Bio-Rad, 161-0301), 0.05% Triton X-100 (Sigma Aldrich, T9284-1L), 0.05% ammonium persulfate (APS, Sigma-Aldrich, A3678), 0.05% tetramethylethylenediamine (TEMED, Sigma-Aldrich, T9281) in DPBS with or without 3 mM benzophenone-methacrylamide (N-[3-[(4-benzoylphenyl)formamido]propyl]methacrylamide, BPMAC PharmAgra). Precursor solutions were prepared as follows: mixture of acrylamide, DI and PBS was degassed *in vacuo* prior to adding SDS, Triton and BPMAC in case of BPMAC+ gel fabrication. Initiators APS and TEMED added immediately prior to addition of mixture to glass slides. The silanized glass slide was sandwiched on top of the wafer shims and precursor mixture was pipetted between the glass slide and the wafer. Excess precursor solution was wicked away with a kimwipe and gels were allowed to polymerize at room temperature.

#### Single-cell SDS-PAGE

EGFP+ HEK-293 cells were lifted with 0.25% trypsin for 10 min and pelleted by centrifugation at 200G for 2 min. Cells were resuspended in 1X DPBS with 1% EDTA (Sigma-Aldrich, 03690-100ML) to prevent cell clustering. 10^6^ cells ml^-1^ were settled into scWestern microwells by gravity and washed to remove excess cells with DPBS. Cell settling was confirmed on a widefield microscope (Nikon Ti2 epifluorescence microscope, motorized stage and filter turret, SOLA SE II 365 light engine, ORCA-Flash4.0 V3 sCMOS camera). The gel was then placed in a custom acrylic chamber with 3D-printed slide-holder insert bounded with platinum wire to enable application of electric field. Cells were lysed with a RIPA-like lysis buffer for 10 seconds, and proteins were then separated by SDS-PAGE at 200V for 40 seconds while imaging by widefield microscopy at 10X magnification.

#### Protein immobilization on nitrocellulose membranes

Protein transfer efficiency to nitrocellulose was quantified over a range of protein target molecular weights. 3 × 0.5 μL of 5 μM EGFP (27 kDa), Ovalbumin Alexa Fluor-555 (46 kDa, ThermoFisher Scientific, O34782), or IgG-Alexa-647 (150 kDa, ThermoFisher Scientific, A-31573) were spotted on 7% BPMAC gel and incubated for 5 min at room temperature in a humidified Petri dish. We then briefly rinsed gel with 1X TBST, imaged the proteins on the gel using widefield microscopy and placed 0.2 μm pore size nitrocellulose membrane on top to transfer protein for 1-4 minutes in triplicate. Next, we imaged polyacrylamide gel to evaluate the fluorescence of the remaining protein. In control experiments, we did not transfer to nitrocellulose but rather allowed the gel to sit for 1-4 minutes to estimate diffusive loss of protein.

#### Evaluation of protein transfer efficiency

We analyzed total fluorescence of protein spots pre- and post-transfer to nitrocellulose. We determined protein mass balance as follows:

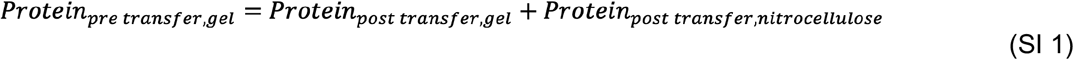

We assumed negligible protein loss to fluid film in the mass balance. We then picked an ROI of identical size for images of pre- and post-transfer that included the outline of the spot and obtained the line-average fluorescence profile. Fluorescence values were adjusted for background by subtracting the lowest fluorescence value in the ROI. We then computed area under the curve for pre- and post-separation images of protein spots and computed efficiency of protein transfer to nitrocellulose as:

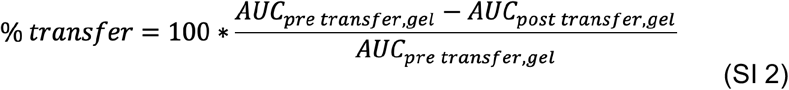

#### Diffusive spreading of immobilized protein

Diffusive spreading was evaluated for scWestern separations immobilized via 1) photocapture in BPMAC-functionalized polyacrylamide or 2) transfer to nitrocellulose from plain polyacrylamide gel. First, EGFP+ HEK-293 cells were settled into 20 μm-diameter wells in 30 μm-thick polyacrylamide gels with and without BPMAC. Cell settling was confirmed by widefield microscopy. Cells were lysed with a RIPA-like lysis buffer^18^ consisting of 0.5% SDS, 0.1% v/v Triton X-100, 0.25% sodium deoxycholate (D6750, Sigma-Aldrich) in 12.5 mM Tris, 96 mM glycine, pH 8.3, (0.5× from a 10× stock, 161-0734, Bio-Rad) for 10 s and separated proteins by SDS-PAGE at 200V for 40 s. Proteins were immobilized by 1) 254 nm UV exposure (Spectrolinker XL 1000 UV) at 3 mW cm^-2^ (Thorlabs PM100D light meter with S120VC 200-1100 nm probe) for 1-4 min on BPMAC-functionalized gel or by 2) transfer to nitrocellulose for 1-4 min. Next, diffusive spreading was quantified by choosing an ROI of identical size for images of pre- and post-immobilization and obtaining line average fluorescence profiles adjusted for background. Fluorescence profiles were plotted and used to compute FWHM by Gaussian fitting in Matlab. To estimate diffusion spreading, FWHM of pre- and post-immobilization protein distributions were analyzed to determine absolute difference and % change. We excluded 3 and 4 min FWHM data for in-gel cases due to low residual fluorescence signal.

#### Mathematical modeling of immobilization kinetics

Reaction-diffusion modeling was performed in Matlab (R2022a, Mathworks). A draft of the code was generated from a text prompt using ChatGPT-3.5 (OpenAI) and manually validated and refined. Partial differential equations were solved on a 1D domain with a spatial resolution of 2 μm for simulation times of 300 s at temporal resolution of 0.02 s. Diffusion constants for EGFP in paper (assumed to be similar to that in water; 89.8 μm^2^ s^-1^) and in 7% polyacrylamide gel (38.1 μm^2^ s^-1^) were estimated using an adjusted Stokes-Einstein diffusivity:

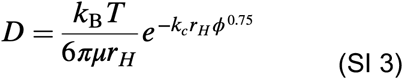

Using hydraulic radius r_H_ = 2.4 nm estimated from the empirical relationship r_H_ = 0.595(M)^0.427^ in nm for globular proteins of molecular weight M (kDa), k_c_ = 0.45 angstrom^-1^, and polymer volume fraction *ϕ* = 0.0093 × %T *−* 0.03151, ref. ^22,23,39–41^. Separate simulations were performed for each Damkohler number, which determined the appropriate value of the immobilization rate constant *k* given a characteristic diffusion length equal to initial scWestern band sizes of ∼100 μm. The initial condition for simulations was a Gaussian band of FWHM = 100 μm with unit amplitude. The PDE describing the unbound fraction was solved using the forward time centered space (FTCS) method and the bound fraction equation was solved using the Euler method. The FWHM of the bound fraction distributions at each time step were estimated by fitting them with Gaussians of standard deviation σ using the nlinfit.m function and determining FWHM = 2σ*√*(2ln(2)).

#### Microwell partitioning fluorescence calibration

To back-calculate in-gel and in-well EGFP concentrations from partition coefficient experiments, calibration was performed by quantifying the fluorescence intensity for a range in EGFP concentrations in a PDMS microwell with channel height of 30 μm (identical to thickness of polyacrylamide gel). We fabricated PDMS microchannels, adhered microchannels to glass slide via plasma bonding and introduced purified EGFP protein labeled with Alexa-555 (ThermoFisher Scientific, A20009) in RIPA-like lysis buffer with 4% BSA (Sigma-Aldrich, A2153-100G) at concentrations of 0.1, 1, 5, 10 μM. We used a confocal microscope to image EGFP fluorescence at the channel mid-plane (Nikon Ti2-E with Yokogawa CSU-W1 spinning disk, 4-line laser source (405nm, 100mW; 488nm, 100mW; 561nm, 100mW; 640nm, 75mW with fiber output), 2048 × 2048 Photometrics Prime BSI sCMOS camera, piezo z stage, and motorized xy stage). We averaged fluorescence over the ROI and then created a calibration curve from concentrations of 0.1, 1, 5, 10 μM by plotting fluorescence vs in-channel concentration of EGFP.

#### Protein partition coefficient measurement

Proteins are expected to partition from hydrogel networks of polyacrylamide gel into free solution, resulting in lower measured in-gel concentration compared to bulk solution^18^. To measure partitioning coefficient, a 7% polyacrylamide gel sandwiched in a microwell hybridization cassette (Arrayit, AHC1X16) was incubated with 40 μL aliquots of 0.1 and 1 μM EGFP labeled with Alexa 555 in RIPA-like lysis buffer with 4% BSA. Gels were allowed to equilibrate for 30 minutes in a covered humidified Petri dish. We then disassembled the hybridization cassette, rinsed the gel with a RIPA-like lysis buffer, sandwiched it with a glass slide to trap protein and quantified in-gel and in-well fluorescence by confocal microscopy. We used the calibration curve we constructed earlier to determine concentration of EGFP in-gel and in wells. We then plotted EGFP concentrations in the gel vs in the well for each tested condition, and used slope of the line to determine partition coefficient (**Figure S1**). Partition coefficient calculated using this method was comparable to previously published values^18^.

#### Single-cell protein detection/LOD using fluorescently-labeled antibody

Fluorescently-labeled antibody detection was compared between EGFP-Alexa-555 tagged protein immobilized in 1) BPMAC-functionalized gel and 2) nitrocellulose post transfer from plain polyacrylamide gel. For both 0.5 μL of EGFP-Alexa-555 concentrations 0.01, 0.0193, 0.0373, 0.072, 0.1389, 0.2683, 0.5179, 1 μM were immobilized in triplicate. On 1) BPMAC-functionalized-gel, the protein dilution series was immobilized by irradiation with 254 nm UVC light for 60 s. Antibody probing was then performed for 2 hours at room temperature with a) 1:200 dilution (0.05 mg ml^-1^) of primary goat polyclonal anti-GFP antibody or b) 1:400 polyclonal rabbit-anti GFP antibody (0.05 mg ml^-1^). We then washed three times with TBST for a total of 1 hour. Next, secondary antibody probing was performed for 1 hr at room temperature with 1:500 dilution (0.004 mg ml^-1^) of a) rabbit anti-goat Alexa-647 tagged antibody or b) donkey anti-rabbit Alexa-647 tagged antibody. Gel was then washed three times with TBST for a total of 1 hour. To evaluate 2) nitrocellulose fluorescently-labeled protein detection,protein dilutions of above mentioned concentrations were first spotted on plain polyacrylamide gel, washed briefly to remove excess, and transferred to nitrocellulose for 3 min. Nitrocellulose was then fixed with 4% PFA for 10 minutes, briefly rinsed with TBST, blocked with 5% BSA in TBST for 30 minutes, and washed three times with TBST for a total of 1 hr and probed with primary and secondary antibodies as described above. Secondary antibody signal was quantified by widefield microscopy at 20X magnification on the Cy5 channel.

Antibody signal-to-noise ratio (SNR) was determined as follows. First, ROI was picked to contain a portion of the inside and edge of the protein spot. Line average fluorescence of the spot adjusted for background was then found to determine signal for the spot. Noise was approximated as average standard deviation of the identical ROI of area near the spot, averaged over concentration conditions to better approximate overall noise. SNR was then calculated as the average fluorescence of the spot divided by noise. Limit of detection was quantified according to Clinical and Laboratory Standards Institute guideline EP17 for limits of detection and quantitation guidelines as follows ^50, 51^:

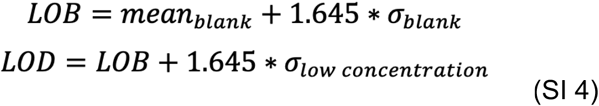

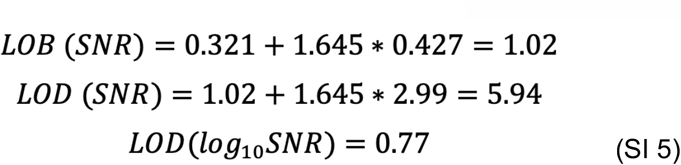

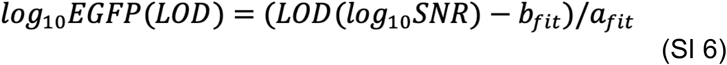

Where *σ*_*blank*_ is standard deviation of the corresponding no protein condition,*σ*_*blank*_ is standard deviation of the lowest EGFP concentration used (0.01 μM), *a*_*fit*_ is slope of the best fit of experiment data plotted as log_10_*SNR* vs log_10_(GFP molecules). For BPMAC probed with Abcam goat anti-GFP antibody, we found *σ*_*blank*_ = 0.42, *σ*_*low concentration*_ = 2.99, *a*_*fit*_ = 0.467, *b*_*fit*_= -2.259, resulting in LOD of log_10_*SNR* = 0.77. The corresponding number of EGFP molecules at LOD is as follows:

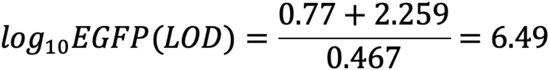

#### Enzyme-linked antibody probing and chemifluorescence detection

EGFP immobilized in nitrocellulose after transfer from plain polyacrylamide gel was used to calibrate the enzyme amplification strategy. 0.5 μL of 1 μM, 100 nM, 10 nM, 1 nM, 100 pM, 10 pM, 1 pM, 100 fM, 0 μM in RIPA-like buffer were spotted in triplicate on 7% BPMAC gel and transferred to nitrocellulose for 3 min. Nitrocellulose was fixed with 4% PFA for 10 min, washed three times with TBST for a total of 60 min, blocked with 5% BSA in TBST for 30 min, and washed three times again in TBST for 60 min total.

Primary antibody probing was performed with 1:200 dilution of goat anti-GFP polyclonal antibody for 2 hr as described above. Nitrocellulose was washed with TBST in triplicate for 60 min. Next, secondary antibody probing was performed with rabbit anti-goat secondary IgG antibody conjugated to HRP for 60 min at room temperature,washed in three times for 60 min with TBST, and twice for 60 min total with TBS. Nitrocellulose was divided into individual protein concentration conditions using a razor blade and placed in a 48-well plate (Corning) for fluorometric detection. QuantaRed™ Enhanced Chemifluorescent HRP Substrate Kit (Thermo Scientific, 15159) was prepared according to the manufacturer specifications. This kit facilitates conversion of ADHP to resorufin in the presence of HRP and H_2_O_2_. 100 μL of HRP substrate was added per well and nitrocellulose was imaged immediately by widefield microscopy via the TRITC channel at 10X magnification and 1 ms exposure every 500 ms for 60 s to quantify increase in resorufin fluorescence. Data were analyzed as follows. An ROI containing the brightest fluorescence within the protein spot was selected. Then the average fluorescence intensity of that ROI was quantified across timepoints. Fluorescence was normalized by the average fluorescence of the no protein control condition for each timepoint to remove background. Normalized average fluorescence was plotted vs time, and slopes were determined over the first 30 s of the timelapse.

To determine the practical detection limit of the enzymatic amplification assay for application in scWesterns, we reasoned that substrate turnover sufficient for successful analyte detection would need to occur within a resorufin reaction product characteristic diffusion time t_D_. This would estimate the time necessary for resorufin to diffuse a characteristic distance ***L*** = 100 μm, or the approximate average diameter of scWestern bands. In other words, analytes can be practically detected only when sufficient substrate turnover occurs to generate a positive signal before diffusion erodes its spatial localization.

We used a published value for resorufin diffusivity in water^52^ and estimated t_D_ as:

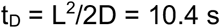

Next we estimate the reaction time t_R_ as the time needed to reach 3.3*S.D. and denoted the practical LOD at the molecule # at which t_R_ < t_D_ transitioned to t_R_ > t_D_. At concentration of 100 pM, t_R_ > t_D_ at 9.5 seconds while at 10 pM we determined that t_R_ < t_D_ at 12 seconds. We then determined an ‘absolute’ LOD regardless of diffusion concerns as the standard deviation of pixel intensities in an ROI for the no-protein control condition at ***t***_***D***_ and computed LOD as:

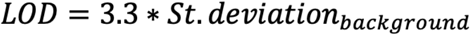

We then compared the delta AFU for calibration protein concentration values at t_D_ ∼ t =10 seconds and determined that EGFP molecule counts above 10 pM concentration (10 ^2.9^ molecules) are above the assay limit of detection yielding dynamic range ∼ 10^3^-10^8^ molecules of EGFP (Figure 3E).

#### Single-cell primary and secondary antibody-based and enzyme-linked antibody protein detection

We tested and compared 1) standard BPMAC in-gel immobilization and probing with primary and secondary fluorescently tagged antibody vs 2) the same for nitrocellulose blotting, 3) enzyme-linked antibody-based EGFP detected in scWestern blots bearing single-cell separations. First, EGFP+ HEK-293 cells were settled into 20 μm-diameter wells in 30 μm-thick polyacrylamide gels with and without BPMAC. Cell settling was confirmed by widefield microscopy and single cells prior to scWestern were imaged on the FITC channel. Cells were lysed with a RIPA-like lysis buffer^18^ for 10 s and separated proteins by SDS-PAGE at 200V for 40 s. Proteins were immobilized by either 1) 1 minute exposure of BPMAC gel to 254 nm UVC or transfer to nitrocellulose for 3 mins for detection schemes 2-4. BPMAC gel washed three times with TBST for a total of 60 min. Nitrocellulose was fixed with 4% PFA for 10 min, washed three times with TBST for a total of 60 min, blocked with 5% BSA in TBST for 30 min, and washed three times again in TBST for 60 min total.

Antibody probing and imaging was performed as follows. For 1) <u>BPMAC-immobilized</u> single-cell proteins, gel was incubated with primary polyclonal goat anti-GFP antibody at 1:20 dilution for 2 hours at room temperature. Gel was then washed with TBST in triplicate for 60 min total. Gel was then incubated with 1:50 dilution of secondary donkey anti-goat Alexa-555 antibody in 1% BSA in TBST for 60 min at room temperature and washed with TBST in triplicate for 60 min total. BPMAC gel was then imaged on widefield TRITC channel at 500 ms exposure at 4X magnification (Nikon). For 2) <u>nitrocellulose-based detection</u>, nitrocellulose was probed with 1:200 dilution for 2 hours at room temperature. Nitrocellulose was then washed with TBST in triplicate for 60 min total. Nitrocellulose was then incubated with 1:500 dilution of secondary donkey anti-goat Alexa-555 antibody for 60 min at room temperature and washed with TBST in triplicate for 60 min total. Nitrocellulose was then imaged on TRITC channel at 500 ms exposure at 4X magnification (Nikon). For 3) <u>Enzyme-linked detection on nitrocellulose</u> in single-cell Westerns, primary antibody probing was performed with 1:200 dilution of goat anti-GFP polyclonal antibody for 2 hours at room temperature. Nitrocellulose was washed with TBST in triplicate for 60 min. Secondary antibody probing was performed with rabbit anti-goat secondary IgG antibody conjugated to HRP for 60 min at room temperature as described above, washed in three times for 60 min with TBST, and twice for 60 min total with TBS. QuantaRed™ Enhanced Chemifluorescent HRP Substrate Kit was prepared according to the manufacturer specifications. 100 μL of QuantaRed kit was added to nitrocellulose containing single-cell Western protein separation, sandwiched with glass slide and the signal was imaged on widefield TRITC channel at 70 ms exposure at 4X magnification (Nikon).

Single cell protein signal was determined as follows. First, we saved circular ROIs for images of single cell EGFP blots immediately after immobilization. We then found the average fluorescence intensity for each ROI. We found background fluorescence as an average of 10 ROIs that do not contain immobilized protein. We then adjusted EGFP fluorescence by background and used it as our final value of initial EGFP fluorescence. Next, we found corresponding ROI for protein signal readout as 1) and 2) Alexa-555 tagged secondary antibody signal, and 3) resorufin signal increase with background subtracted. We then plotted initial EGFP signal vs readout and created a fit. LOD was determined as 3.3*S.D of the blank.

## Supplementary figures

**Figure S1.**
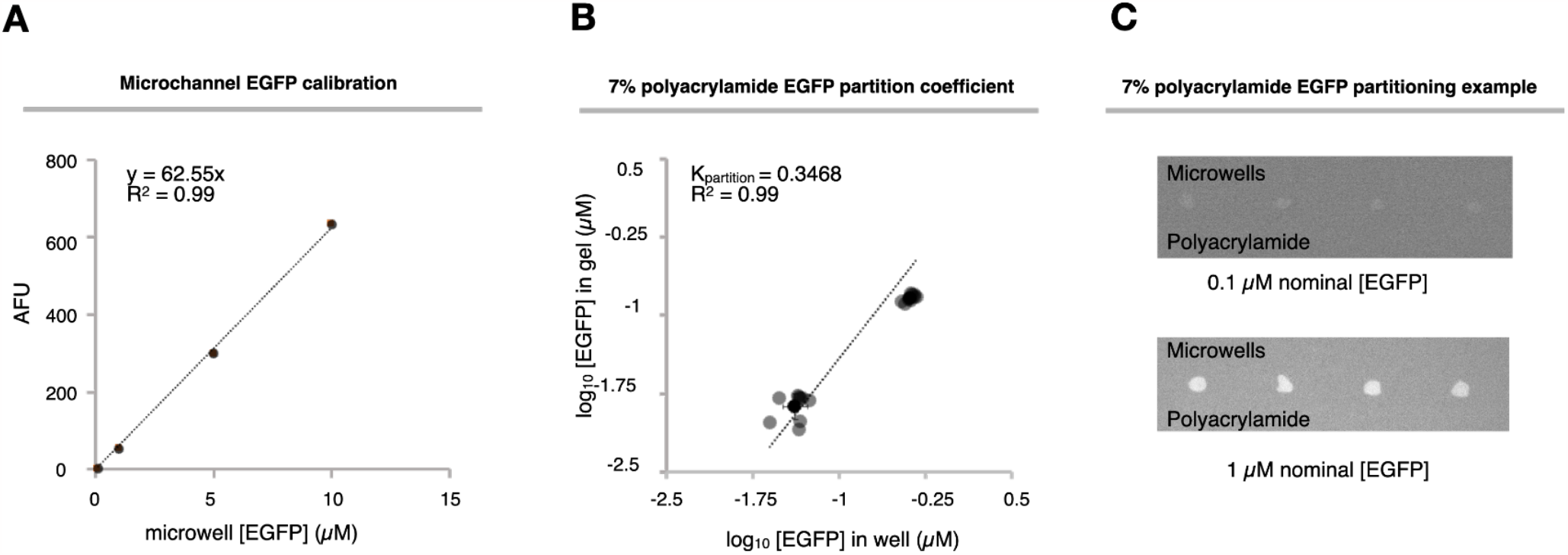
EGFP microwell concentration calibration curve and determination of EGFP partition coefficient in 7%T polyacrylamide gels. (**A**) Calibration curve for EGFP labeled with AlexaFluor 555 in denaturing RIPA buffer in PDMS microchannels of 30 μm height as measured on a confocal microscope. EGFP concentrations of 0.1, 1, 5 and 10 μM were converted into a number of molecules per characteristic size of scWestern band 100 μm in diameter and 30 μm tall. (**B**) Partition coefficients were determined for several select EGFP-Alexa Fluor 555 concentrations in the RIPA-like buffer including 0.1 and 1 μm, with *n* = 3 for each concentration. 7%T polyacrylamide gel was incubated with EGFP-Alexa Fluor 555 concentration in hybridization cassette wells as 40 μL aliquots for 30 mins, rinsed, protein trapped with glass slide and in-gel and in-well fluorescence measured on a confocal microscope. Calibration curve was used to determine in-gel and in-well concentrations of EGFP-AlexaFluor 555, concentrations plotted as log_10_ and partition coefficient was determined as slope of the resultant line. (**C**) Fluorescence micrographs of EGFP partitioning in microwells and in polyacrylamide gel at equilibrium for varying EGFP concentrations. Contrast settings are equivalent.

**Figure S2.**
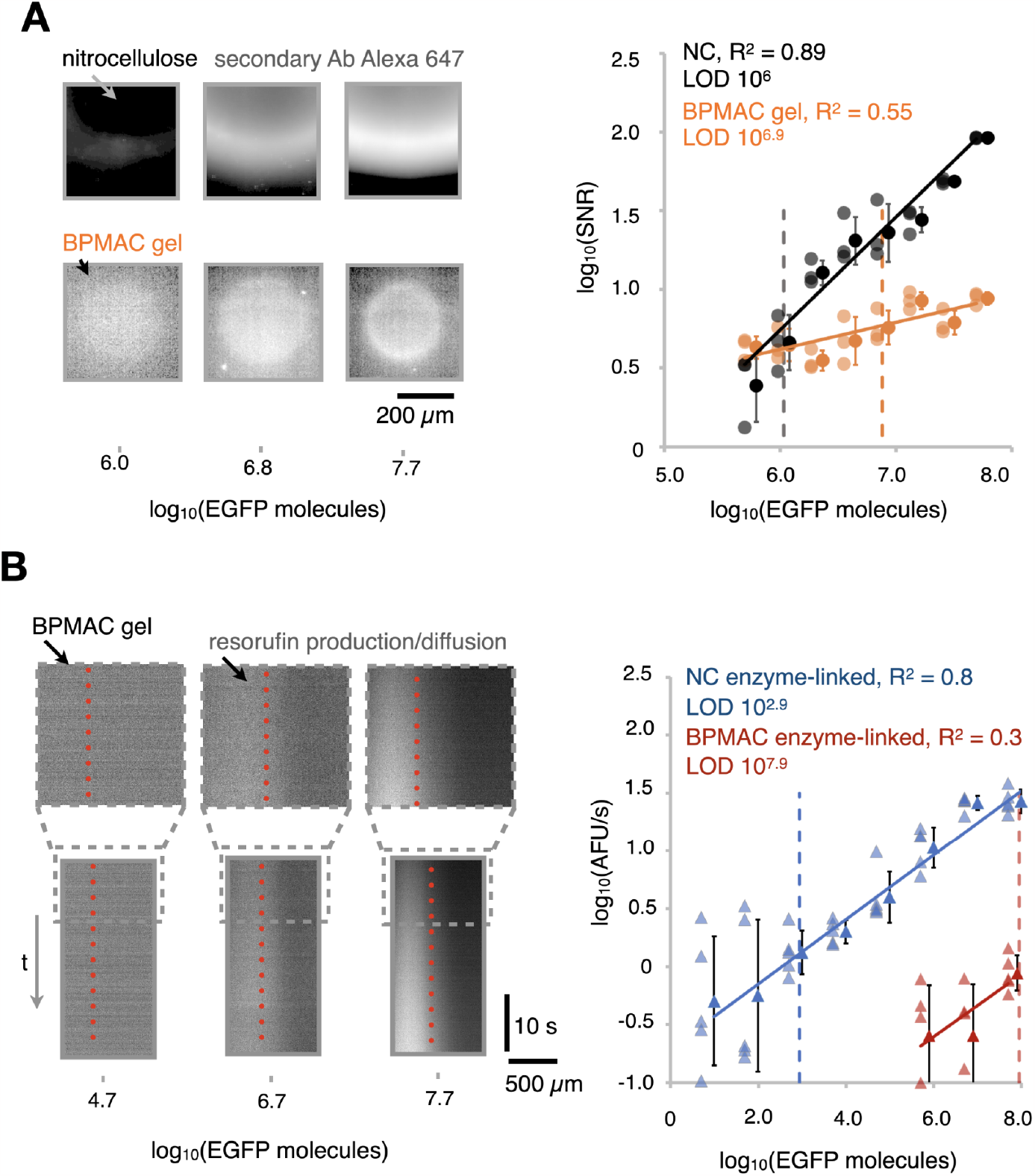
Additional calibration data for fluorescently-labeled and enzyme-linked antibody detection strategies. **(A)** *Left*, immunofluorescence micrographs of calibration spots of EGFP transferred to nitrocellulose or immobilized in-gel prior to probing with Invitrogen rabbit anti-EGFP primary and anti-rabbit Alexa-647 secondary antibody, pixel intensity was log-transformed. Note that contrast settings are equivalent within each image row, but different between rows. *Right*, corresponding calibration curves. (**B**) *Left*, immunofluorescence kymographs of calibration spots of EGFP immobilized in 7% BPMAC gel and detected with HRP-conjugated secondary antibodies for enzymatic amplification. Note that concentration of 1 nM corresponding to log_10_(EGFP molecules) = 4.7 had no appreciable increase in fluorescence compared to background. Contrast settings are equivalent within image row. *Right*, corresponding calibration curves for HRP-based protein detection on nitrocellulose (blue) and on BPMAC gels (red).

